# Injectable hydrogels with phase-separated structures that can encapsulate live cells

**DOI:** 10.1101/2022.01.31.478579

**Authors:** Shohei Ishikawa, Yuki Yoshikawa, Hiroyuki Kamata, Ung-il Chung, Takamasa Sakai

**Affiliations:** Department of Bioengineering, School of Engineering, The University of Tokyo, 7-3-1 Hongo, Bunkyo-ku, Tokyo, Japan; Center for Disease Biology and Integrative Medicine, School of Medicine, The University of Tokyo, 7-3-1 Hongo, Bunkyo-ku, Tokyo, Japan; Department of Materials Engineering, School of Engineering, The University of Tokyo, 7-3-1 Hongo, Bunkyo-ku, Tokyo, Japan

**Author notes:** These authors contributed equally to this work (Shohei Ishikawa, Yuki Yoshikawa, Hiroyuki Kamata). Corresponding authors: (H.K.), (T.S.).

## Abstract

Injectable hydrogels are biomaterials that can be administered minimally invasively in liquid form, and are considered promising artificial extracellular matrix (ECM) materials. However, ordinary injectable hydrogels are synthesized from water-soluble molecules to ensure injectability, resulting in nonphase-separated structures, making them structurally different from natural ECMs with phase-separated insoluble structural proteins, such as collagen and elastin. Here, we propose a simple material design approach to impart phase-separated structures to injectable hydrogels by adding inorganic salts. Injecting a gelling solution of mutually crosslinkable tetra-arm poly(ethylene glycol)s with potassium sulfate at optimal concentrations results in the formation of a hydrogel with phase-separated structures in situ. These phase-separated structures provide up to an 8-fold increase in fracture toughness while allowing the encapsulation of live mouse chondrogenic cells without compromising their proliferative activity. Our findings highlight that the concentration of inorganic salts is an important design parameter in injectable hydrogels for artificial ECMs.

## Main Text

Hydrogels, consisting of water and a network of three-dimensionally crosslinked polymers, are promising biomaterials that function as an artificial extracellular matrix (ECM) due to their high cytocompatibility and mass diffusion properties.^1,2^ Injectable hydrogels, which are synthesized by injecting crosslinkable polymer components with a syringe,^3^ and more recently, three-dimensional bioprinters,^4^ are of particular importance in terms of securing minimally invasive properties and material formability. Despite their potential biomedical applications, the scope of application of ordinary injectable hydrogels remains limited because of their nonphase-separated or nonporous structure,^5^ consequently appearing transparent in color.^6^ By contrast, natural biological tissues or ECMs have rather opaque phase-separated structures, as the precursors of structural proteins such as collagen^7^ and elastin^8^ are insolubilized by complex posttranslational processing, forming porous structures,^9^ thus providing mechanical toughness^10^ and space for cellular activity.^11^ The lack of phase-separated structures in ordinary injectable hydrogels stems from the fact that they rely on the crosslinking of molecules that remain water-soluble throughout the synthesis process to ensure injectability. Intuitively, the use of water-soluble molecules is contradictory to the production of insoluble phase-separated structures. Therefore, it is necessary to introduce a temporally dynamic process in which initially water-soluble molecules become insoluble over time, thereby forming phase-separated structures in situ.

Poly(ethylene glycol) (PEG), which is often used in the synthesis of artificial ECMs due to its high biocompatibility,^2,12^ is known to undergo phase separation in aqueous solutions containing certain molecules such as polysaccharides^13^ and inorganic salts.^14^ In particular, this tendency is more pronounced for larger molecular weights (*M*_w_) and higher concentrations of PEG (*C*_PEG_). Taking advantage of such properties and noting that *M*_w_ always increases over time during the formation of hydrogels, Broguiere et al. combined crosslinkable PEG and dextran to achieve temporally dynamic phase separation in hydrogels.^15^ However, the addition of polysaccharides is not an ideal practice due to the following reasons: (i) unknown contaminants cannot be ruled out as being of biological origin, (ii) different manufacturing methods result in different physical properties, thus requiring fine-tuning to ensure reproducibility across different laboratories,^16^ and (iii) the bioinert nature of PEG is compromised. Therefore, it is undoubtedly beneficial to produce phase-separated structures by combining a single synthetic polymer species (i.e., PEG only) with inorganic salts, which, to our knowledge, have never been applied to injectable hydrogels.

Here, we hypothesized that PEG, which is initially water-soluble, would spontaneously undergo phase separation when crosslinked in the presence of inorganic salts at an appropriate concentration above a certain threshold (*C*_critical_). Furthermore, we assumed that a balance between the timing of phase separation and gelation would result in a PEG hydrogel that inherited the characteristics of phase-separated structures. To test this hypothesis, injectable hydrogels were synthesized from mutually crosslinkable tetra-arm PEGs (Tetra-PEG) using the concentration of inorganic salts at the material preparation stage (*C*_IS_) as a key parameter. We showed that injectable PEG hydrogels with phase-separated structures could be obtained using potassium sulfate (K_2_SO_4_) at optimal concentrations. Our elongation test results demonstrated that the phase-separated structures improved the toughness of the obtained hydrogels. We also showed that live mouse chondrogenic cells (ATDC5) could be encapsulated in the interconnected pore structure without loss of their proliferative activity. The addition of inorganic salts to existing injectable hydrogels is expected to further expand research related to the development of artificial ECMs.

## Results

### Design of injectable hydrogels with phase-separated structures

Prior to the design of hydrogels, we dissolved hydroxy-terminated linear PEGs (Linear-PEG) of various *M*_w_ in aqueous solutions with and without K_2_SO_4_ to reproduce the known static phase separation (**Fig. 1a**). We observed that Linear-PEGs with *M*_w_ ≤ 10 kg/mol were completely dissolved regardless of the presence of K_2_SO_4_, resulting in transparent aqueous solutions. In contrast, we noticed that the aqueous solutions of Linear-PEGs with *M*_w_ ≥ 100 kg/mol became opaque in the presence of K_2_SO_4_, forming a partial precipitate in the case of the Linear-PEG with the highest *M*_w_ = 1000 kg/mol. This was also evident in the absorbance measurement (**Fig. 1b**). Taking this static phenomenon as a hint, we devised a synthetic scheme in which mutually crosslinkable Tetra-PEGs are crosslinked to increase *M*_w_ over time in aqueous solutions containing K_2_SO_4_ so that the growing PEG network spontaneously undergoes phase separation beyond a certain time point to provide hydrogels with phase-separated structures (**Fig. 1c**). Here, we used a combination of sulfhydryl- and succinimidyl-terminated Tetra-PEGs (Tetra-PEG-SH and Tetra-PEG-OSu, respectively) to form thioester crosslinks.

**Fig. 1.**
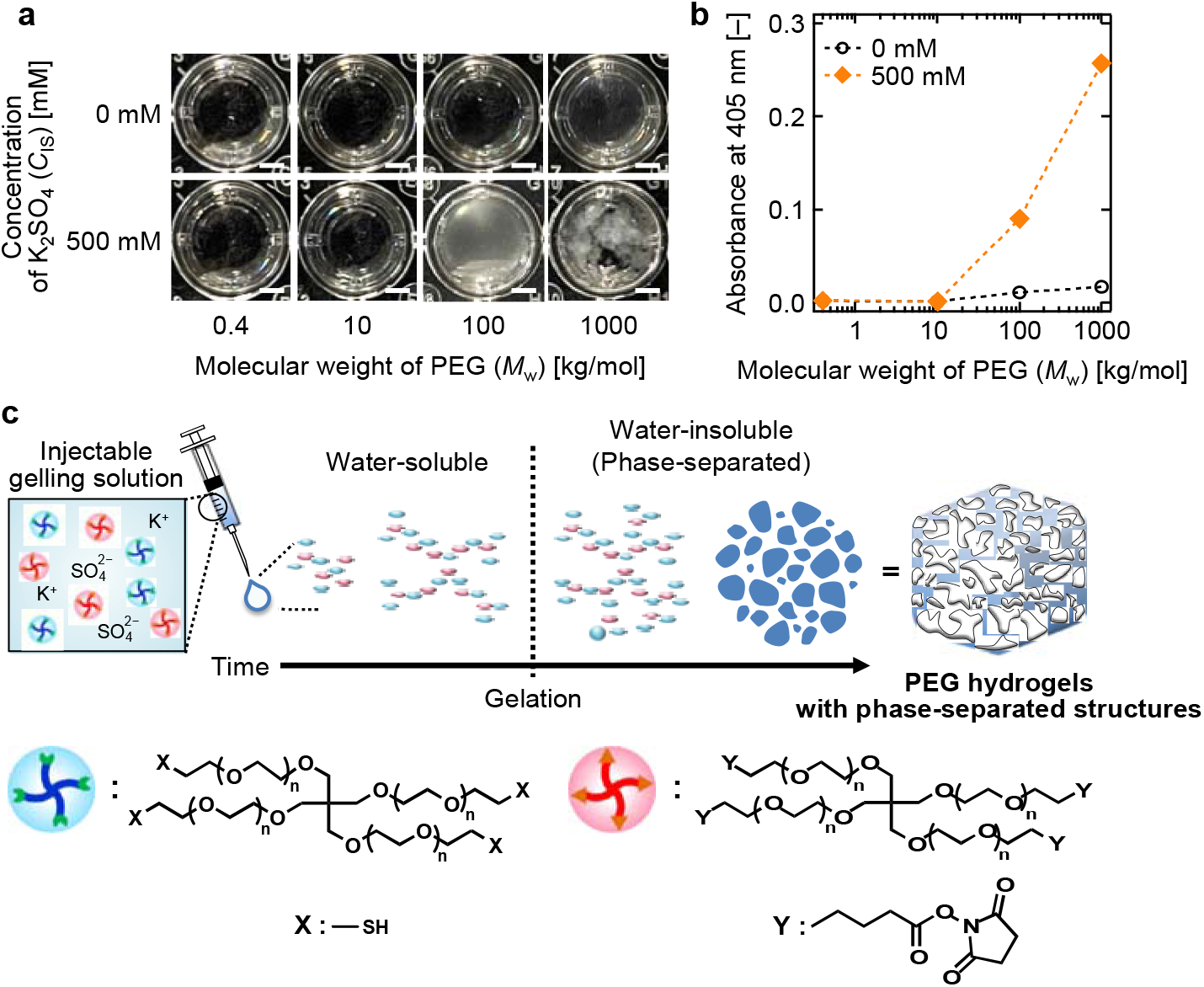
Conceptual design of injectable hydrogels with phase-separated structures. **a** Images of aqueous solutions of Linear-PEGs with various *M*_w_, *C*_PEG_ = 50 g/L, and *C*_IS_ = 0 mM (top) or 500 mM (bottom). Scale bars represent 2 mm. **b** Absorbance at 405 nm of aqueous solutions of Linear-PEGs with various *M*_w_, *C*_PEG_ = 50 g/L, and *C*_IS_ = 0 mM (open circle) or 500 mM (orange diamond). Dashed lines between symbols are shown as guide. **c** Schematic illustration showing the temporally dynamic phase separation of PEG caused by K_2_SO_4_ present at *C*_IS_ ≥ *C*_critical_, which results in the formation of PEG hydrogels with phase-separated structures.

### Formation and characterization of hydrogels

Based on the theoretical design, we synthesized hydrogels by dissolving Tetra-PEG-SH and Tetra-PEG-OSu in phosphate buffered aqueous solutions at pH 7.4 in the presence or absence of K_2_SO_4_, and then mixed equal volumes of the 2 PEG solutions. First, we conducted the gelation test using K_2_SO_4_ (**Fig. 2a**). We found that the aqueous solutions of Tetra-PEG-SH and Tetra-PEG-OSu (hereafter referred to as precursor solutions) with *M*_w_ = 10 kg/mol and *C*_PEG_ = 100 g/L were transparent even in the presence of K_2_SO_4_. The liquid prepared by mixing the 2 precursor solutions (hereafter referred to as gelling solution) was also transparent immediately after mixing. It is important to note that the gelling solution containing K_2_SO_4_ could be injected through a 27G needle with an outer diameter = 0.22 ± 0.03. When we observed the injected gelling solution after 5 min, we noticed that the liquid had lost its fluidity, forming an opaque solid. In this study, we defined the phenomenon of loss of macroscopic fluidity when tilting the vial as gelation, while the time required to induce gelation was defined as gelation time (*t*_gel_). We accordingly observed that the appearance of hydrogels after gelation differed markedly between those with and without K_2_SO_4_ (**Fig. 2b**). More specifically, in the absence of K_2_SO_4_, the mean value of *t*_gel_ was 5.3 min, and the hydrogel remained transparent even beyond *t*_gel_. Whereas, in the presence of K_2_SO_4_, the hydrogel formed in a similar way and the mean value of *t*_gel_ was 3.6 min, but became opaque after *t*_gel_. We also observed this opacity in the same materials in absorbance measurements, in which we noticed a sharp increase in absorbance after *t*_gel_ only in the presence of K_2_SO_4_ (**Fig. 2c**). We also performed the gelation test by adding a red fluorescent dye to the gelling solution, and observed the obtained structure using confocal laser scanning microscopy (CLSM) (**Fig. 2d**). We found that in the absence of K_2_SO_4_, no characteristic structure was apparent either before or after gelation. In contrast, in the presence of K_2_SO_4_, we noticed a phase-separated sea-island structure beyond *t*_gel_, with the size of pores being on the order of 10^-6^ to 10^-5^ m. Interestingly, this sea-island structure did not develop further from the point of formation. Image reconstruction from the data obtained by scanning the z-axis showed that the sea-island structure was formed in 3 dimensions (**Fig. 2e**). When we immersed the formed hydrogels in India ink, an example of colloids with particle size of 10^-7^ to 10^-5^ m,^17^ we noticed that the black color did not penetrate into the hydrogel without a phase-separated structure, whereas it penetrated into the interior of hydrogels with phase-separated structures (**Fig. 2f**). We also conducted gelation tests using gelling solutions with various *C*_PEG_ and *C*_IS_, and characterized their appearance by visual observations (**Fig. 2g**), CLSM images (**Fig. 2h**) (see **Supplementary Figure 1**) and absorbance measurements (**Fig. 2i**). We observed a trend in which high *C*_PEG_ resulted in low *C*_critical_. Briefly, 200 mM ≤ *C*_critical_ ≤ 250 mM at *C*_PEG_ = 20 g/L, and 150 mM ≤ *C*_critical_ ≤ 200 mM at *C*_PEG_ ≥ 50 g/L. Even in the region of *C*_IS_ ≥ *C*_critical_, we detected that the microstructure was sensitive to *C*_IS_; especially at *C*_PEG_ = 100 g/L, the sea-island structure was more pronounced at *C*_IS_ = 250 mM than at 200 mM, and furthermore, at *C*_IS_ = 300 mM, we observed a particle-like cocontinuous structure, which slightly increased transparency.

**Fig. 2.**
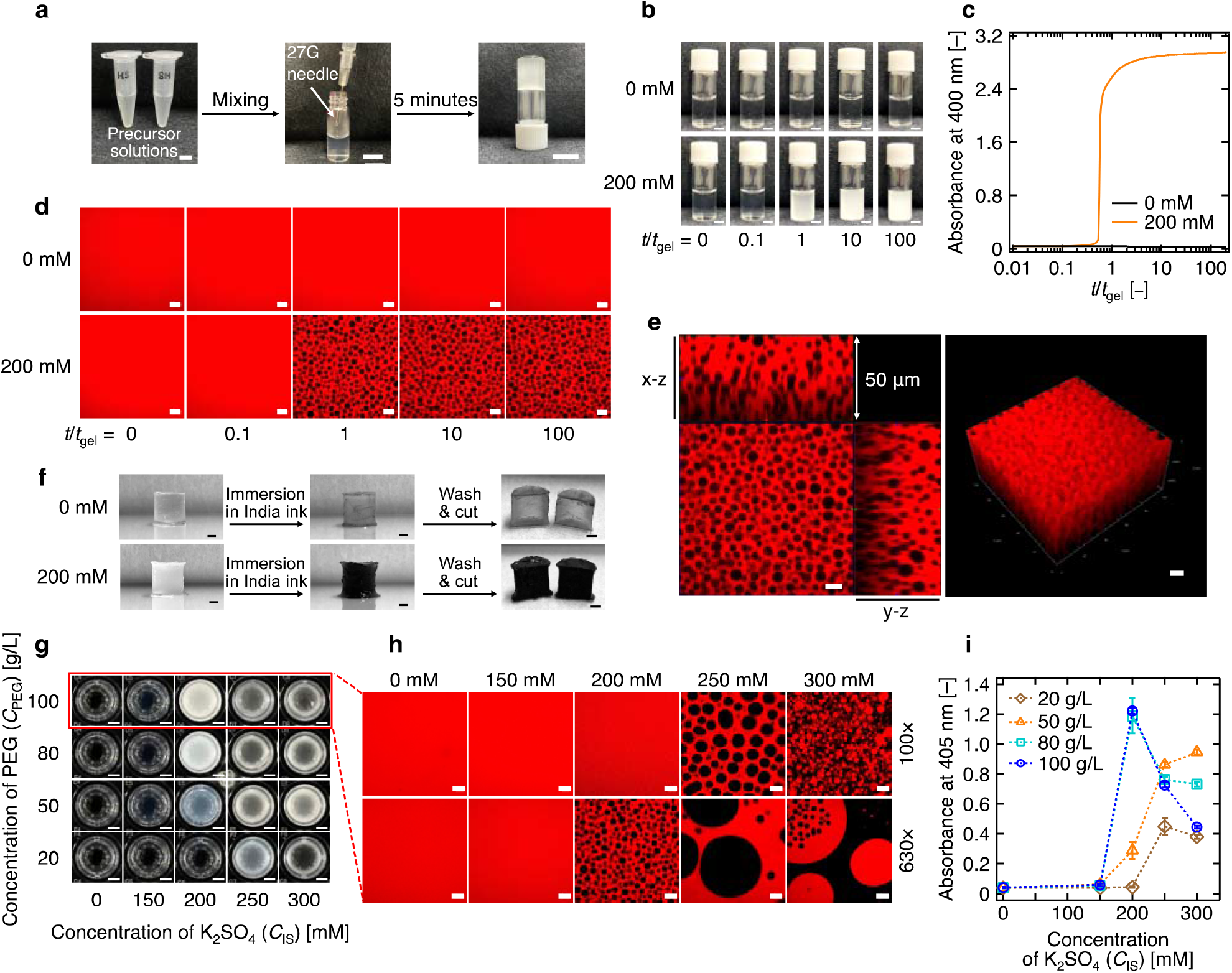
Characterization of the formation process of hydrogels and the formed hydrogels. **a** Hydrogel formation process. Equal volumes of precursor solutions with *C*_PEG_ = 100 g/L and *C*_IS_ = 200 mM (left) are mixed generating the gelling solution which is then injected via a 27G syringe needle (middle) to form the hydrogel (5 min after injection; right). Scale bars represent 10 mm. **b** Images of gelling solutions with *C*_PEG_ = 100 g/L and *C*_IS_ = 0 mM (top) or 200 mM (bottom) over time. Time elapsed since the preparation of gelling solutions (*t*) is normalized by *t*_gel_. Scale bars represent 5 mm. **c** Time course of absorbance at 400 nm of gelling solutions with *C*_PEG_ = 100 g/L and *C*_IS_ = 0 mM (black line) or 200 mM (orange line). **d** CLSM images of gelling solutions or formed hydrogels with *C*_IS_ = 0 mM (top) or 200 mM (bottom) during the gelation process. Scale bars represent 10 μm. **e** The x-z and y-z planes of the CLSM image obtained for the hydrogel with *C*_IS_ = 200 mM at *t*/*t*_gel_ = 100, aligned with the x-y plane (left). Three-dimensional image of the formed hydrogel with *C*_IS_ = 200 mM reconstructed from CLSM images at *t*/*t*_gel_ = 100 (right). Scale bars represent 10 μm. **f** Images of hydrogels prepared with *C*_PEG_ = 100 g/L and *C*_IS_ = 0 mM (top) or 200 mM (bottom) after being immersed in India ink. Scale bars represent 2 mm. **g** Images of hydrogels prepared with various *C*_PEG_ as a function of *C*_IS_. Scale bars represent 2 mm. **h** CLSM images of hydrogels prepared with various *C*_IS_ and *C*_PEG_ = 100 g/L. Scale bars represent 50 μm and 10 μm for 100× and 630× magnification, respectively. **i** Absorbance at 405 nm of hydrogels prepared with *C*_PEG_ = 20 (brown diamond), 50 (orange triangle), 80 (turquoise square), and 100 g/L (blue circle) as a function of *C*_IS_. Error bars represent the standard deviation of the mean obtained from 2 samples. Dashed lines between symbols are shown as guide.

### Evaluation of aqueous stability of phase-separated structures

We examined the aqueous stability of phase-separated structures of hydrogels prepared in a cylinder mold with various *C*_IS_. In this study, we defined the as-prepared state as the state immediately after removal from the mold. We immersed the hydrogels in the as-prepared state in DPBS and allowed them to equilibrate until the degree of swelling (*Q*) reached a plateau; we specifically defined *Q* at this point as the equilibrium degree of swelling (*Q*_eq_). We found that in the as-prepared state, hydrogels with *C*_IS_ ≥ 200 mM were opaque (**Fig. 3a**). Although we recognized an increase in *Q*_eq_ that was dependent on *C*_IS_ (see **Supplementary Figure 2**), hydrogels did not become completely transparent and did not match the appearance of hydrogels prepared without K_2_SO_4_ even at equilibrium (**Fig. 3b**). CLSM images showed that the phase-separated structures generated for *C*_IS_ ≥ 200 mM were microscopically maintained at equilibrium with a negligible increase in their pore size.

**Fig. 3.**
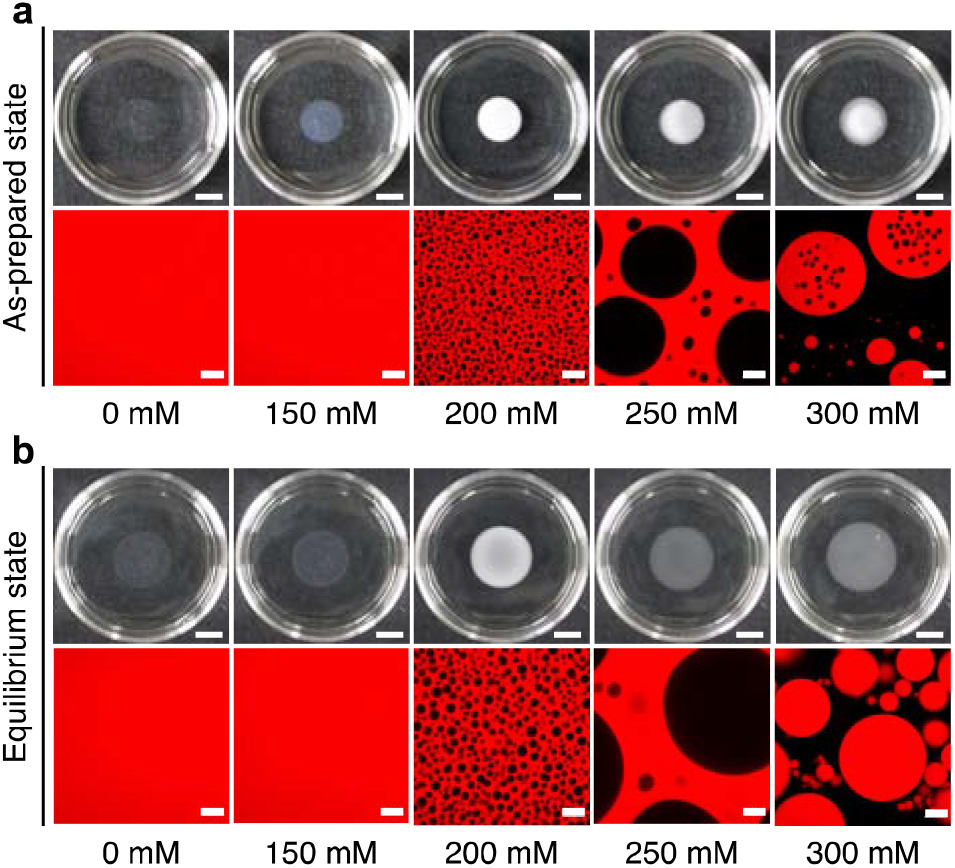
Stability of the phase-separated structures in hydrogels in an aqueous environment. **a** Optical (top) and CLSM (bottom) images of hydrogels in the as-prepared state with various *C*_IS_ and *C*_PEG_ = 100 g/L. Scale bars represent 10 mm and 20 μm for optical and CLSM images, respectively. **b** Optical (top) and CLSM (bottom) images of hydrogels in the equilibrium state with various *C*_IS_ and *C*_PEG_ = 100 g/L. Scale bars represent 10 mm and 20 μm for optical and CLSM images, respectively.

### Mechanical characterization of hydrogels

To investigate the effect of phase-separated structures on the mechanical properties of the resulting hydrogels, we prepared hydrogels with various *C*_IS_, and analyzed their mechanical properties using elongation tests. We did not perform any elongation test on the hydrogel with *C*_IS_ = 300 mM as it was too soft to be elongated. We noticed that the representative stress-elongation curves of hydrogels clearly differed depending on *C*_IS_ (**Fig. 4a**). In addition, the stress at fracture (*σ*_max_) showed a gradual increase depending on *C*_IS_. Interestingly, we found that for *C*_IS_ = 200 mM, *σ*_max_ was exceptionally high, more than 2-fold higher than that for *C*_IS_ = 0 mM (**Fig. 4b**). We observed a similar discontinuous trend in the elongation ratio at fracture (*λ*_max_), which was specifically high at *C*_IS_ = 200 mM and approximately 3-fold higher than that for *C*_IS_ = 0 mM (**Fig. 4c**). We further noticed that as *σ*_max_ and *λ*_max_ increased, the fracture energy, which is used as an index of material toughness, also showed a similar trend (**Fig. 4d**). The Young’s modulus (*E*) was around 20 kPa for *C*_IS_ ≤ 200 mM, whereas it was as high as 30 kPa for *C*_IS_ = 250 mM (**Fig. 4e**).

**Fig. 4.**
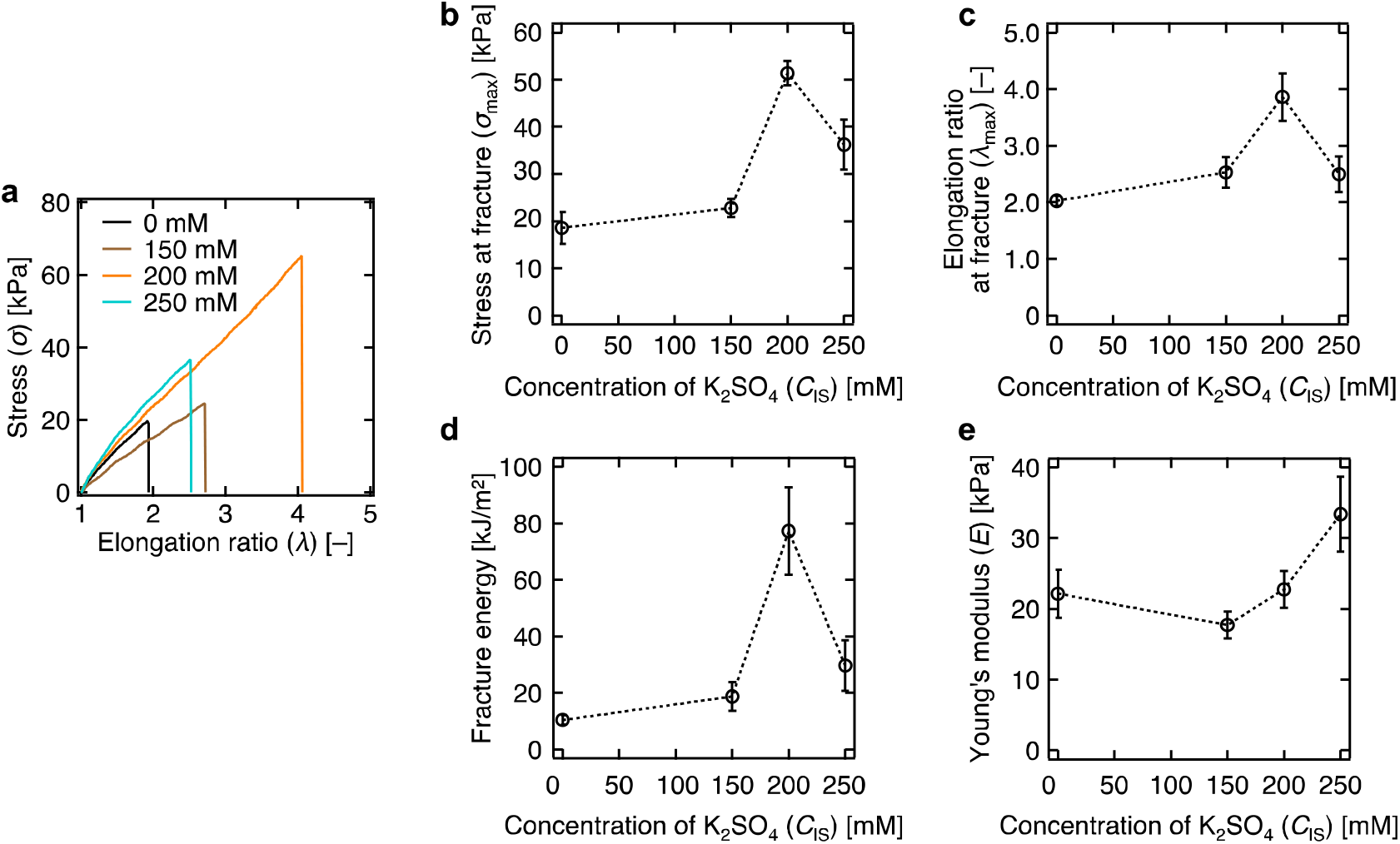
Mechanical properties of hydrogels. **a** Stress-elongation curves for hydrogels prepared with *C*_PEG_ = 100 g/L and *C*_IS_ = 0 (black line), 150 (brown line), 200 (orange line), and 250 mM (turquoise line). Only one representative example from each group is depicted. **b** *σ*_max_, **c** *λ*_max_, **d** fracture energy, and **e** *E* of hydrogels prepared with *C*_PEG_ = 100 g/L and various *C*_IS_. Error bars represent the standard deviation of the mean obtained from 5 samples. Dashed lines between symbols are shown as guide.

### In vitro cell encapsulation studies

To evaluate the cytocompatibility of phase-separated structures generated in a time-dependent manner, we prepared hydrogels using presuspended live cells in gelling solution and inducing gelation in that state. Here, we prepared hydrogels containing live ATDC5 cells using hydrogels with various *C*_IS_. As a representative example, we prepared a hydrogel with *C*_IS_ = 200 mM and performed a viability assay of the encapsulated ATDC5 cells (**Fig. 5a**), in which live and dead cells were detected as green and red signals, respectively. We qualitatively confirmed the uniform dispersion of green signals. We calculated the cell viability immediately after encapsulation in both cases, either with or without K_2_SO_4_ (**Fig. 5b**). In another experiment, we encapsulated ATDC5 cells in hydrogels with various *C*_IS_, and then stained the polymer component of hydrogels, filamentous actin (F-actin), and nuclei of cells using red, green, and blue fluorescent dyes, respectively (**Fig. 5c**). We detected differences in the dispersion of the red signal depending on *C*_IS_, corresponding to the previous observation without ATDC5 cells. We noticed that the pattern of the green signal was also *C*_IS_-dependent; when *C*_IS_ = 0 mM, the green signal was again observed to be uniformly dispersed in the field of view, but the number of green signals tended to decrease with an increase in *C*_IS_; this trend was in good agreement with the pattern of the blue signal. Focusing on the composite images of the red, green, and blue signals, we clearly noticed that the cell-derived green and blue signals were localized in the voids created by phase separation. To confirm that the proliferative activity of ATDC5 cells encapsulated in hydrogels was not altered, we collected ATDC5 cells from hydrogels with phase-separated structures and examined their subsequent proliferation behavior. First, to confirm the reported degradability of hydrogels with thioester crosslinks, we immersed a hydrogel prepared with *C*_IS_ = 200 mM in an aqueous solution containing _L_-cysteine methyl ester hydrochloride (_L_-cys) (see **Supplementary Figure 3**). We observed that in the _L_-cys-free environment, *Q* reached a plateau after a time-dependent increase, whereas in the _L_-cys-containing environment, *Q* was increased continuously, accompanied by a slight increase in pore size, and the hydrogel was eventually completely dissolved. We then encapsulated ATDC5 cells in the hydrogel with *C*_IS_ = 200 mM. Subsequently, the hydrogel was degraded with _L_-cys without conducting cell culture, and we retrieved the ATDC5 cells in the liquid and seeded them onto twodimensional cell culture plates for culturing. We compared the growth of the retrieved ATDC5 cells with that of ATDC5 cells that did not undergo the process of encapsulation in hydrogels. Using the same cell viability assay as in the earlier experiment, we observed the lack of any morphological differences between the groups tested (**Fig. 5d**). Similarly, when we stained F-actin and cell nuclei using fluorescent dyes (red and purple, respectively), again we did not observe any obvious differences between the groups (**Fig. 5e**). We further confirmed this result by quantitatively evaluating the growth rate of cells from the same culture experiments (**Fig. 5f**).

**Fig. 5.**
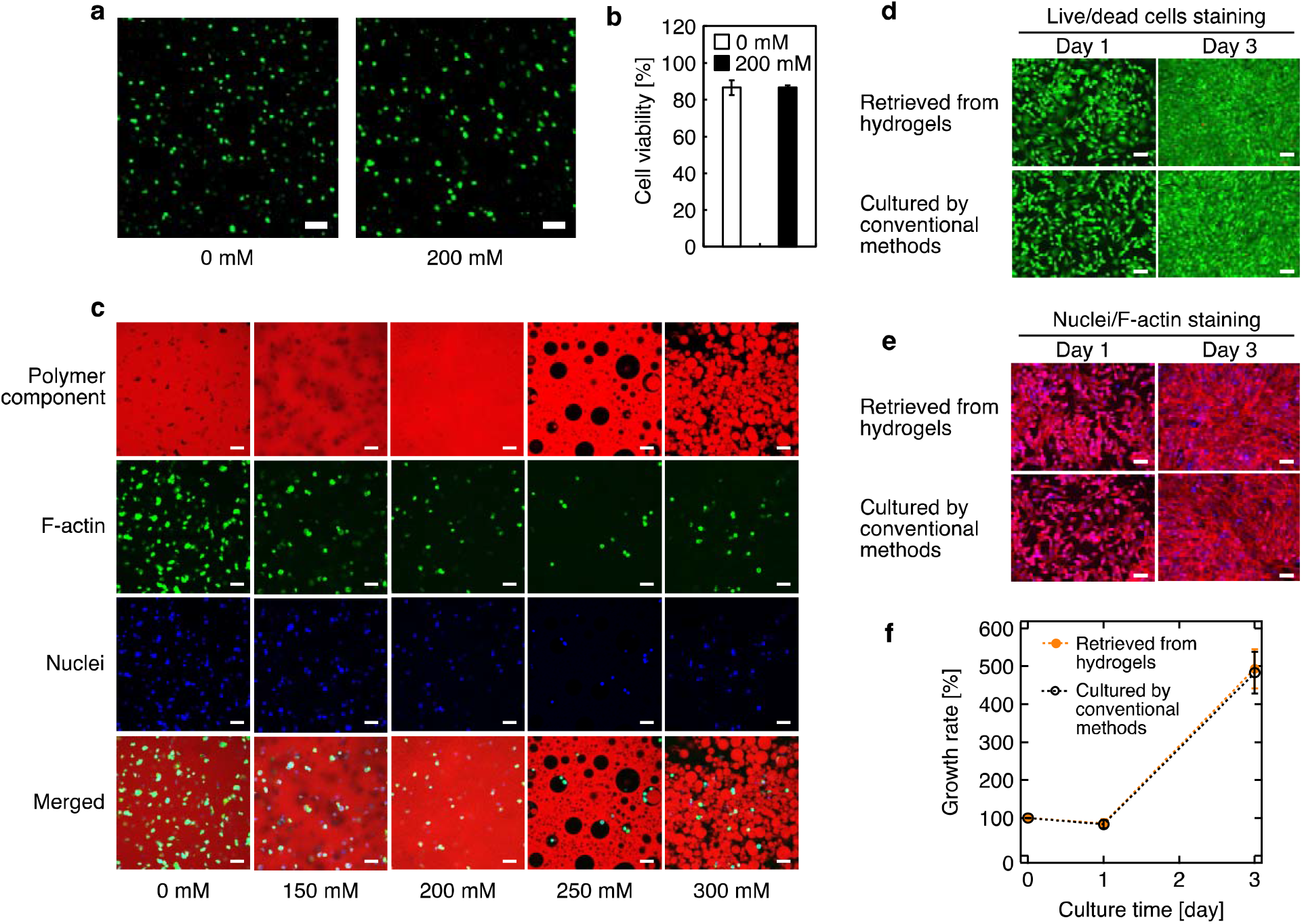
In vitro evaluation of cytocompatibility. **a** Fluorescence microscopy images of hydrogels formed from gelling solutions with various *C*_IS_ containing ATDC5 cells, where green and red signals represent live and dead cells, respectively. Scale bars represent 50 μm. **b** Cell viability calculated from fluorescence images. Error bars represent the standard deviation of the mean obtained from 3 samples. **c** Fluorescence microscopy images of hydrogels formed from gelling solutions with various *C*_IS_ containing ATDC5 cells, where red, green, and blue signals represent polymer components, Factin, and cell nuclei, respectively. Scale bars represent 50 μm. Comparison of cell activity between **d** the group of ATDC5 cells once encapsulated in a hydrogel and then retrieved and recultured on a cell culture plate, and **e** the control group of ATDC5 cells (nonencapsulated in hydrogels). Green and red signals represent live and dead cells, respectively. For F-actin and cell nuclei, red and purple signals were used, respectively. Scale bars represent 200 μm. **f** Growth rate in two-dimensional culture of ATDC5 cells once encapsulated in a hydrogel and then retrieved (orange circle), and nonencapsulated ATDC5 cells (open circle). Error bars represent the standard deviation of the mean obtained from 3 samples. Dashed lines between symbols are shown as guide.

## Discussion

In this study, we tested the hypothesis that phase-separated structures can be easily obtained by adding inorganic salts at optimal concentrations during the synthesis of injectable PEG hydrogels. We chose PEG as the target polymer for phase separation because of its high biocompatibility and accumulated knowledge of its use as an ECM substrate.^12^ Among the inorganic salts that are known to cause the phase separation of PEG, we chose K_2_SO_4_, which is classified as a neutral salt, to take advantage of its documented effect on the physical properties of PEG,^18^ and to avoid pH changes, which might affect gelation time^19^ and cellular activities.^20^ Potassium salts are also thought to have another advantage in that the ionic species do not interfere with the sodium phosphate buffer system used to adjust pH for controlling *t*_gel_.^9^ However, from the standpoint of cell physiology, the use of sodium ions would be preferable outside the cell, and such sodium salts (e.g., sodium sulfate) would be the ideal choice. For the synthesis of hydrogels, we used mutually crosslinkable Tetra-PEGs, which require only 2 essential components, water and PEG,^21,22^ to which we simply added K_2_SO_4_ as a phase separation inducer. Although a number of hydrogels have been synthesized using branched PEGs with various end-groups and their ECM applications have been reported,^23^ we chose a combination of sulfhydryl and succinimidyl end-groups to form thioester crosslinks because of their advantages of being usable at biologically relevant pH, allowing the three-dimensional encapsulation and on demand _L_-cys-induced safe retrieval of live cells.^24^

The phase separation observed in aqueous PEG solutions was successfully reproduced by K_2_SO_4_ in the crosslinking process of PEG, resulting in opaque hydrogels at *C*_IS_ ≥ *C*_critical_. It is worth noting that the phase-separated structures were introduced in a time-dependent manner, allowing hydrogels to be treated as liquid before injection, and subsequently inducing their phase separation in situ. We noticed that *t*_gel_ was shorter in the presence of K_2_SO_4_ due to the accelerated crosslinking in the gel phase that held the condensed precursors. The phase-separated structures obtained exhibited sea-island structures, which were microscopically similar to those commonly observed in polymer blends.^25^ Considering the components included in the present system, it is safe to say that the observed sea-island structures were simply composed of PEG that became hydrophobic and phase-separated around *t*_gel_ due to the involvement of K_2_SO_4_. The z-stack measurements of the phase-separated structures and the colloid immersion test indicated that the generated porous structures were not partial but interconnected in 3 dimensions. Such interconnected structures are essential for the design of artificial ECMs because they facilitate the interaction of the three-dimensionally hydrogel-encapsulated cells with the outside environment,^26^ and allow space for cellular activities.^27^ Furthermore, the fact that the interconnected pores were observed even in centimeter-sized hydrogels suggested that they are scalable in size and can be useful for prosthetics for large defects in the body.

The qualitative trend of *C*_critical_ to be lower at higher *C*_PEG_ was consistent with the results of the liquid experiments conducted here and in previous studies.^14^ Interestingly, even above *C*_critical_, not only the on-off boundary of phase separation was determined, but also the microstructure was altered. More specifically, to obtain the desired microstructure, it is necessary to select the appropriate *C*_IS_. Although the change in the *C*_IS_ sensitive microstructure is an interesting phenomenon, we did not include it in the scope of this study, as it is a topic that requires extensive research. As predicted by previous studies, the optimal *C*_IS_ was different for each *C*_PEG_.^14^ This finding indicated that the optimal *C*_IS_ needs to be carefully selected when changing *C*_PEG_, for example, to adjust the mechanical properties of the hydrogel.

Unlike experiments using hydrogels in the as-prepared state, ECMs are continuously exposed to excess aqueous body fluids (e.g., extracellular fluid).^28^ Therefore, we considered it important to investigate whether the phase-separated structures observed in the as-prepared hydrogels would be maintained in their equilibrium state. Swelling tests showed that the phase-separated structures were stable in an aqueous environment, suggesting the possibility of their use as an ECM. This behavior was consistent with that of PEG hydrogels with phase-separated structures prepared using noninjectable approaches.^29^

The establishment of phase-separated structures is known to affect the resulting mechanical properties.^30,31^ Therefore, we investigated whether the same effects could be observed in the resulting hydrogels. Overall, we observed that the physical properties of hydrogels showed a gradual increase dependent on *C*_IS_. This was assumed to be the result of 2 opposing aspects caused by phase separation: (i) condensation of PEG in the gel phase and (ii) decrease in the volume fraction of the gel phase. Improvement in these mechanical properties suggested that the effect of the former outweighed the latter, in consistency with the nonlinear polymer concentration dependence of *σ*_max_, *λ*_max_, and *E* in polymer hydrogels^32^. The *C*_PEG_ at the time of measurement was not characterized in this study due to technical issues regarding the dependence of the time of expulsion of water on the size of hydrogels^33^ and its loss by evaporation. Even if those effects were considered, the hydrogel prepared with *C*_IS_ = 200 mM was anomalously resistant to fracture compared with the groups prepared with other *C*_IS_ values; although this unique property was lost at much higher *C*_IS_ values. These results indicated that the phase separation occurring in hydrogels must not be excessive if toughness is desired, and that the concentration of inorganic salts must be carefully adjusted to obtain sea-island structures, such as those observed in the hydrogel with *C*_IS_ = 200 mM; that is, it must not be adjusted to form a particle-like cocontinuous structure as seen in *C*_IS_ = 300 mM. In contrast, the situation was different for *E*, as we did not find any singular point in the range we tested. It is unclear at this stage whether the lack of a singular point is a universal fact or simply could not be observed. Comparing the hydrogel with the phase-separated structure at *C*_IS_ = 200 mM with the control hydrogel prepared without K_2_SO_4_, the fracture energy was improved approximately 8-fold. This is direct evidence that the introduction of inorganic salts adds toughness to injectable PEG hydrogels.

In light of the fact that this technique can also be applied to other injectable PEG hydrogels, the focus should be on the ratio of improvement rather than each absolute value. A limitation of the current study is that it misses the concept of relaxation originating from the porous structure.^34^ Therefore, further studies need to examine the viscoelastic properties in more detail to gain a better understanding.

K_2_SO_4_ induced phase-separated structures in hydrogels even in the presence of cell culture-related reagents, such as culture medium and live ATDC5 cells. The phase-separated structures showed similar morphological trends regardless of the presence or absence of cells, indicating that the proposed method can be applied for the design of ECMs. Addition of inorganic salts to a cell culture system is known to increase osmotic pressure (i.e., hyperosmotic environment), which might cause osmotic shock to live cells.^28,35^ However, in the present study, the addition of K_2_SO_4_ did not significantly increase cell death, thus making it a useful approach for cell culture applications. Although we used ATDC5 cells in this study as a representative example of live cells, other cell types might also be used for encapsulation, as shown in previous studies.^36^ For hydrogels with phase-separated structures, the number of ATDC5 cells that could be encapsulated in hydrogels was reduced. This was attributable to the fact that water was drained out due to phase separation, potentially carrying away cells out of the hydrogel. We are confident that our results demonstrated the effective encapsulation of cells in phase-separated structures; however, this quantitative issue of cell loss needs to be addressed in future studies. We did not perform long-term cultures because of the well-known effect of anoikis (i.e., apoptosis upon loss of attachment to ECMs) in unmodified PEG hydrogels.^37^ Similarly, the effect of phase-separated structures on cell behavior was not fully investigated, as the optimal pore structure is known to vary with each cell type.^38^ Therefore, it would be worthwhile to apply this method to existing nonphase-separated but multifunctional PEG hydrogels and experiment with different cell types and physiological stimuli.

Broadly, these results showed that the addition of inorganic salts to injectable PEG hydrogels can easily produce phase-separated structures with interconnected pores that contribute to toughness and the encapsulation of live cells. Because of the simplicity of the method, it might be applicable to existing nonphase-separating hydrogels and could stimulate the research and development of artificial ECMs.

## Methods

### Materials

Sulfhydryl-terminated tetra-arm poly(ethylene glycol) (PEG), also known as PTE-100SH, and succinimidyl-terminated tetra-arm PEG, also known as PTE-100HS, with *M*_w_ = 10 kg/mol (Tetra-PEG-SH and Tetra-PEG-OSu, respectively) were purchased from NOF CORPORATION (Tokyo, Japan). Potassium sulfate (K_2_SO_4_), sucrose, 4% paraformaldehyde phosphate buffer solution (PFA), dimethyl sulfoxide (DMSO), and _L_-cysteine methyl ester hydrochloride (_L_-cys) were purchased from FUJIFILM Wako Pure Chemical Corporation (Tokyo, Japan). Hydroxy-terminated linear PEGs (Linear-PEG) with *M*_w_ = 0.4, 10, 100, and 1000 kg/mol were purchased from Sigma-Aldrich (Missouri, USA). Alexa Fluor™ 594 C_5_ Maleimide (Alexa-MA), Alexa Fluor™ 594 Phalloidin (Alexa-Ph), LIVE/DEAD™ Viability/Cytotoxicity Kit, *-Cellstain*®-Hoechst 33258 solution (Hoechst 33258), phosphate-buffered saline (PBS), Dulbecco’s phosphate-buffered saline (DPBS), Dulbecco’s modified Eagle’s medium (DMEM), penicillin/streptomycin (PS), trypsin-EDTA (0.05%), 0.4% trypan blue solution, and fetal bovine serum (FBS) were purchased from Thermo Fisher Scientific (Massachusetts, USA). India ink was purchased from KAIMEI & Co., Ltd. (Saitama, Japan). Alexa-MA was prepared as a 1 mg/mL DMSO solution. Phosphate buffer (pH 7.4) (PB) was prepared at a concentration of 200 mM as previously reported.^19^ The RC15-AC 0.22 *μ* m filter (Sartorius AG, Göttingen, Germany), ARVO™ X3 microplate reader (PerkinElmer, Inc., Massachusetts, USA), LSM 800 confocal laser scanning microscope (CLSM) (Carl Zeiss AG, Jena, Germany), and M165C optical microscope (Leica Camera AG, Wetzlar, Germany) were used in all experiments. Milli-Q water was used throughout the study.

### Phase separation and absorbance measurement of linear PEG

Linear PEGs with *M*_w_ = 0.4, 10, 100, and 1000 kg/mol were separately dissolved in PB to give a concentration of PEG (*C*_PEG_) = 50 g/L. After 24 h, solutions were filtered through a 0.22 μm filter. K_2_SO_4_ was added to filtered solutions to obtain aqueous solutions with a concentration of K_2_SO_4_ (*C*_IS_) = 0 or 500 mM. Then, 300 μL of each liquid was transferred to a 96-well plate and images were captured under an optical microscope. Absorbance of each well was then measured at 405 nm using a microplate reader.

### Observation of gelation and absorbance measurement of formed hydrogels

Tetra-PEG-SH and Tetra-PEG-OSu were separately dissolved in PB with *C*_IS_ = 0-300 mM to obtain *C*_PEG_ = 20-100 g/L that were termed precursor solutions. Precursor solutions with the same *C*_PEG_ and *C*_IS_ were mixed in equal volumes (gelling solution) and poured into a vial, mold, or well in 96-well plates using a micropipette or syringe. Subsequently, the gelling solution was incubated at 25°C for 24 h to obtain hydrogels. For 96-well plates, 200 μL was poured in each well. Absorbance of hydrogels formed in a 96-well plate was measured at 405 nm using a microplate reader. Images were captured at each time point during the process.

### Absorbance measurements during gelation process

A gelling solution was poured into a plastic cuvette with an optical path length of 10 mm. Absorbance at 400 nm was measured using a UV-vis spectrophotometer, V-670 (JASCO Corporation, Tokyo, Japan) at 25°C every 5 s for 24 h.

### Microscopic observation of phase-separated structures

Tetra-PEG-SH was dissolved in PB with *C*_IS_ = 0-300 mM to obtain *C*_PEG_ = 100 g/L. Alexa-MA was added to the solution to 1 vol%, and the mixture was incubated at 25°C for 10 min. Similarly, Tetra-PEG-OSu was dissolved in PB with the same *C*_PEG_ and *C*_IS_. Equal volumes of the 2 precursor solutions were mixed, poured into a cylindrical silicone mold (diameter: 5 mm and height: 1 mm), and incubated at 25°C for 24 h. Obtained hydrogels were observed under CLSM. For 3D observation, z-axis scans were obtained with an interval of 0.35 μm.

### India ink immersion tests

A gelling solution with *C*_PEG_ = 100 g/L and *C*_IS_ = 0 or 200 mM was poured into a cylindrical mold made of polytetrafluoroethylene (PTFE) (diameter: 7 mm and height: 7 mm) and incubated at 25°C for 24 h. Each hydrogel was carefully removed from the mold and immersed in India ink. Hydrogels were pushed several times with a finger, and then rinsed with excess water and split in 2 parts. Images were captured during the process.

### Swelling tests and calculation of degree of swelling

Hydrogels with *C*_PEG_ = 100 g/L and *C*_IS_ = 0 to 300 mM were prepared in a PTFE mold (diameter: 15 mm, height: 7 mm) and then removed from the mold. This state was defined as-prepared state. Hydrogels were immersed in DPBS and incubated at 25°C for 24 h to reach their equilibrium state. The diameters of the 2 states *(d_0_* and *d*_eq_, respectively) were measured using an optical microscope, and the degree of swelling in the equilibrium state (*Q*_eq_) was calculated as *Q*_eq_ = (*d*_eq_/*d*_0_)^3^, assuming that hydrogels deform isotropically. CLSM images of hydrogels in the as-prepared and equilibrium states were captured.

### Elongation tests and evaluation of physical properties

Uniaxial elongation tests were conducted using a tensile testing apparatus, Autograph AG-X plus (Shimadzu Corporation, Kyoto, Japan). A load cell of 10 N was used in all tests. A gelling solution with *C*_PEG_ = 100 g/L and *C*_IS_ = 0 or 250 mM was poured into a dumbbell-shaped mold (JIS K6251:2017 Standards) made of silicone and incubated at 25°C for 24 h. Each hydrogel was carefully removed from the mold and elongated at a head speed of 60 mm/min while recording the load. Stress-elongation (*σ-λ*) curves were calculated from the obtained load and displacement data. For each hydrogel composition, the same test was performed at least 5 times.

### Encapsulation of live cells

Live mouse chondrogenic cells (ATDC5) (passage 19, 1.0 × 10^6^ cells) were expanded in 100 mm × 20 mm dishes (Corning Inc., New York, USA) using 10 mL DMEM containing 10% FBS and 1% PS and cultured at 37°C under 5% CO_2_ for 1 week. During cell culture, the medium was replaced every 2-3 d. Before the encapsulation of ATDC5 cells into hydrogels, Tetra-PEG-SH and Tetra-PEG-OSu solutions with *C*_PEG_ = 100 g/L and *C*_IS_ = 0 or 200 mM containing 10% sucrose were sterilized using a 0.22 μm filter. Alexa-MA was added to the Tetra-PEG-SH solution to 1 vol%, and the mixture was incubated at 25°C for 10 min. ATDC5 cells (1.0 × 10^5^ cells) were dispersed in 10 μL Tetra-PEG-SH solution in a 1.5 mL tube. This cell suspension was mixed with an equal volume of Tetra-PEG-OSu solution. The mixture was then incubated for 15 min at 37°C under 5% CO_2_ to obtain hydrogels. The final cell density in the hydrogel was 5.0 × 10^6^ cells/mL.

### Cell viability assay for cells encapsulated in hydrogels

For this assay, we used the LIVE/DEAD™ Viability/Cytotoxicity Kit and prepared 500 μL DMEM containing 1 μM calcein AM (for live cells; green) and 2 μM ethidium homodimer (for dead cells; red) according to the manufacturer’s instructions. Hydrogels containing ATDC5 cells were immersed in the prepared DMEM, stained over 10 min, and then fixed with PFA for 10 min in the dark under 5% CO_2_ at 37°C. Afterwards, hydrogels were washed 3 times with DPBS before being observed under a CLSM. The number of live and dead cells was counted using ImageJ software and cell viability was calculated as the ratio of live cells to total cells.

### Triple staining of polymer component, F-actin, and cell nuclei

Hoechst 33258 was diluted in DMEM to a concentration of 5 μg/mL. Hydrogels containing ATDC5 cells were immersed in 500 μL of the prepared Hoechst 33258 solution and incubated in the dark for 1 h. Then, hydrogels were immersed in 500 μL PFA for 20 min and further incubated in 500 μL 0.5% Triton™ X-100 for 5 min. Hydrogels were additionally incubated in 1% Alexa-Ph for 3 h before being observed under a CLSM. All staining steps were performed at 37°C. Between each step, hydrogels were washed thrice with PBS.

### Degradation test

A gelling solution was poured into a cylindrical glass mold (diameter: 1.9 mm) and incubated at 25°C for 24 h before being removed from the mold. Here, PB was diluted 2-fold with water to obtain 100 mM PB (pH 7.4) into which _L_-cys was dissolved to a concentration of 5 mM (degradation buffer). Hydrogels were immersed in 30 mL degradation buffer at 25°C and changes in color and diameter were observed under an optical microscope over time. Following the procedures as in other microscopic observation, CLSM images of degrading hydrogels were captured.

### Evaluation of cellular proliferative activity

Hydrogels containing ATDC5 cells were immersed in degradation buffer containing 10% sucrose and incubated at 37°C under 5% CO_2_ for 60 min. The mixture was centrifuged at 1000 rpm for 3 min and the supernatant liquid was removed.

Retrieved ATDC5 cells were suspended in DMEM, and seeded onto 12-well plates at 1.0 × 10^5^ cells/well. As a control, the same number of nonencapsulated ATDC5 cells were also seeded in 12-well plates. Cell viability assay and staining for cell nuclei and F-actin were performed at 1 and 3 d after seeding. For the cell viability assay, calcein AM and ethidium homodimer were used as per the manufacturer’s instructions before observations by a CLSM. Similarly, Hoechst 33258 and Alexa-Ph were used for staining cell nuclei and F-actin according to the manufacturer’s instructions. For the evaluation of growth rate, ATDC5 cells were detached from culture plates by adding 500 μL trypsin-EDTA (0.05%) and incubating for 5 min at 37°C under 5% CO_2_. Detached cells were collected and centrifuged at 1000 rpm for 3 min. Following the removal of the supernatant, 500 μL DMEM was added and cells were stained using trypan blue. Stained cells were then counted using an automated cell counter (Thermo Fisher Scientific, Massachusetts, USA).

## Acknowledgements

This work was supported by the Japan Society for the Promotion of Science (JSPS) (grant number: 20J01344) and Japan Agency for Medical Research and Development (AMED) (grant number: JP20im0210821).

## Author contributions

H.K., T.S., and U.C. designed the study. Y.Y, S.I., and H.K. designed the experiments. Data acquisition and/or analysis was performed by S.I and Y.Y. H.K. and S.I. drafted the manuscript. Administrative, technical, or supervisory tasks were handled by T.S., U.C., and H.K.

---

**Supplementary Figure 1.**
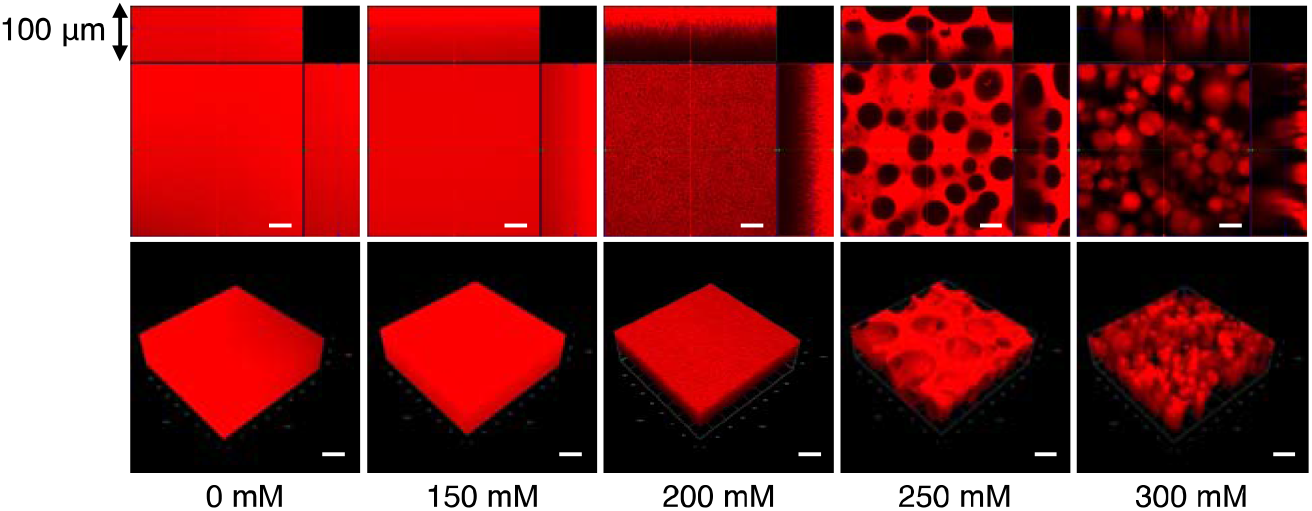
Three-dimensional images of hydrogels with various *C*_IS_ and *C*_PEG_ = 100 g/L reconstructed from CLSM images. Scale bars represent 40 μm.

**Supplementary Figure 2.**
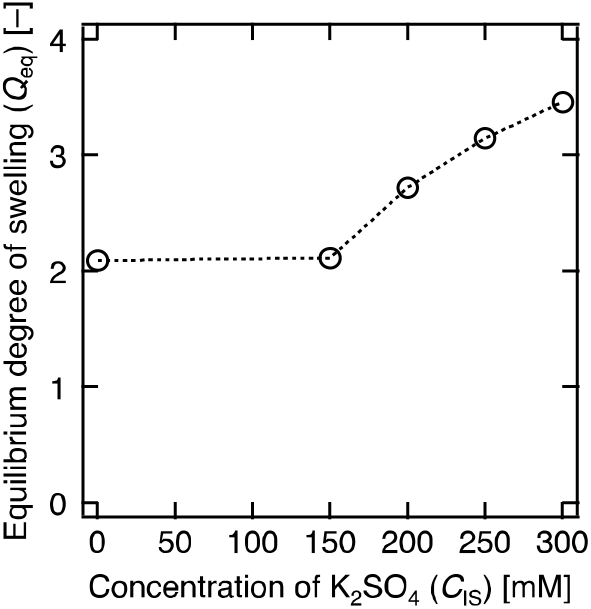
*Q*_eq_ of hydrogels prepared with various *C*_IS_ and *C*_PEG_ = 100 g/L. Dashed lines between symbols are shown as guide.

**Supplementary Figure 3.**
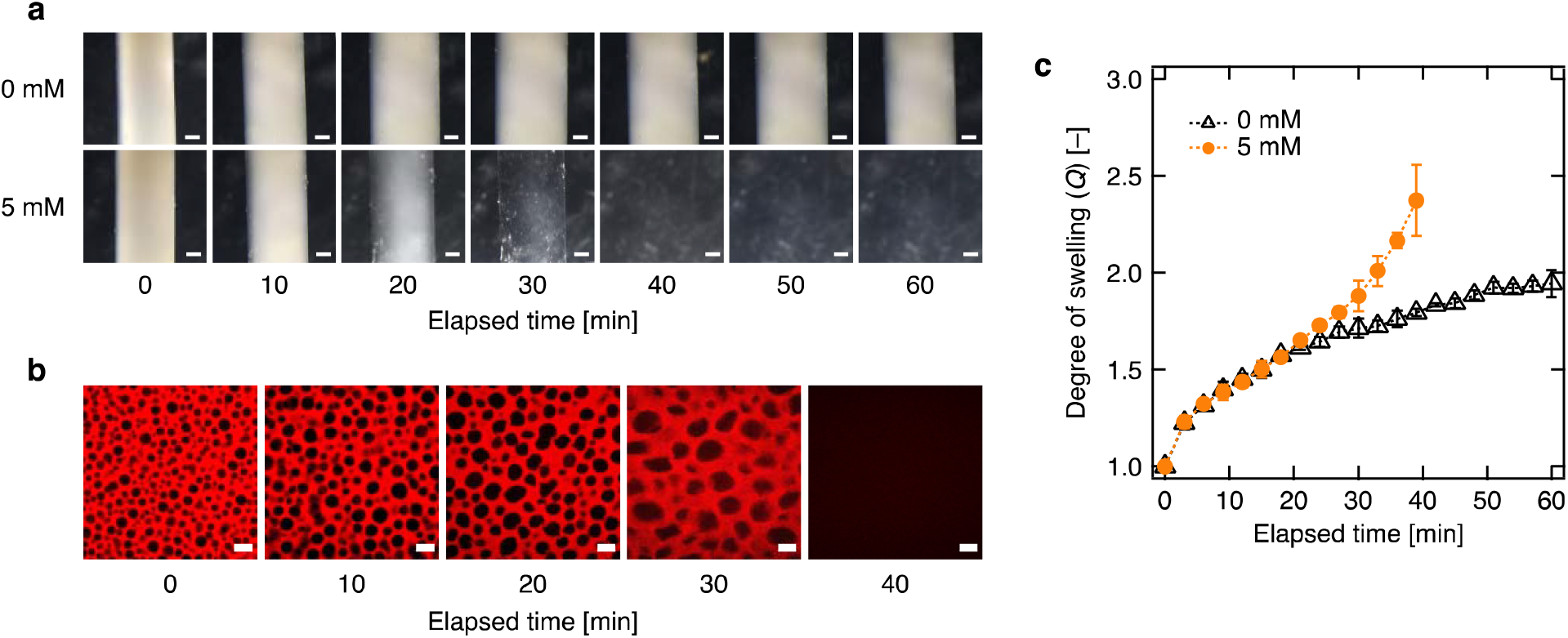
Degradation tests of hydrogels prepared with *C*_PEG_ = 100 g/L and *C*_IS_ = 200 mM. **a** Time course images of hydrogels immersed in aqueous solutions containing _L_-cys at 0 mM (top) and 5 mM (bottom). Scale bars represent 500 μm. **b** CLSM images of degrading hydrogels. Scale bars represent 10 μm. **c** Time variation of *Q* for hydrogels immersed in aqueous solutions containing _L_-cys at 0 mM (open triangle) and 5 mM (orange circle). Error bars represent the standard deviation of the mean obtained from 2 samples. Dashed lines between symbols are shown as guide.

